# Manipulating patterns of dynamic deformation elicits the impression of cloth with varying stiffness

**DOI:** 10.1101/556530

**Authors:** Wenyan Bi, Peiran Jin, Hendrikje Nienborg, Bei Xiao

## Abstract

Cloth is a common material and humans can visually estimate its mechanical properties by observing how it deforms under external forces. Here we ask whether, and how, dynamic deformation can affect the perception of mechanical properties of cloth. In Experiment 1, we find that both intrinsic mechanical properties and optical properties affect stiffness perception when the stimuli are presented as images. By contrast, in videos, humans can partially discount the effect of optical appearances and exhibit higher sensitivity to stiffness. We further identified an idiosyncratic deformation pattern (i.e., movement uniformity) to differentiate stiffness, which can be reliably measured by six optical flow features. In Experiment 2, we isolate the deformation by creating dynamic dot stimuli from the 3D mesh of the cloth. We directly alter the movement pattern by manipulating the uniformity of the displacement vectors on the dot stimuli, and show that changing the pattern of dynamic deformation alone can alter the perceived stiffness of cloth in a variety of scene set-ups. Furthermore, by analyzing optical flow fields extracted from the manipulated dynamic dot stimuli, we confirmed the same six optical flow features can be diagnostic of the degree of stiffness of moving cloth across different scenes. Overall, our study demonstrates that manipulating patterns of dynamic deformation alone can elicit the impression of cloth with varying stiffness, suggesting that human visual system might rely on the idiosyncratic pattern of dynamic deformation for estimating stiffness.

## Introduction

For humans to successfully navigate in the environment, it is crucial to estimate material properties of objects and predict how they would react under external forces. In the last two decades, a good amount of measurements have been made on how humans perceive material properties of rigid objects (e.g., surface gloss of a plastic object) (Bi, Newport, & Xiao, 2018; Fleming, Gegenfurtner, & Nishida, 2015; Fleming, 2017; Maloney & Brainard, 2010). Numerous image features have been proposed for the perception of optical properties of solid objects (Fleming, Wiebel, & Gegenfurtner, 2013; Kim, Marlow, & Anderson, 2012; Motoyoshi, 2010; Motoyoshi, Nishida, Sharan, & Adelson, 2007; Sawayama & Nishida, 2018). However, many objects around us are non-rigid and deformable, such as cloth, food, and liquids. They deform and move under externally applied forces. In this situation, seeing how they deform over time would be informative about their mechanical properties such as soft or hard, heavy or light, liquid or solid, elastic or stiff (Paulun, Schmidt, Assen, & Fleming, 2017).

One way to quantify the dynamic deformation is to measure the movement pattern through analyzing features of the optical flow fields extracted from the videos. Previous research suggests that humans could use distortions in the optical flow fields to estimate mechanical properties (e.g., softness) of an object (Bi, Jin, Nienborg, & Xiao, 2018; Kawabe, Maruya, Fleming, & Nishida, 2015; Nishida, Kawabe, Sawayama, & Fukiage, 2018; Paulun et al., 2017; Van Assen, Barla, & Fleming, 2018). However, visually estimating mechanical properties of deformable objects is challenging because several factors including optical properties (e.g., gloss, transparency), intrinsic mechanical properties (e.g., malleability), as well as external forces can affect the perceived deformation pattern (see Figure 1) and cause distortions in the optic flow conjointly. For example, the deformation pattern generated by crumpling a skirt is very different from that of a flapping flag under wind force. Hence, if the goal is to estimate the intrinsic mechanical properties of deformable objects, the visual system has to disentangle the causal contributions of these factors.

**Figure 1:**
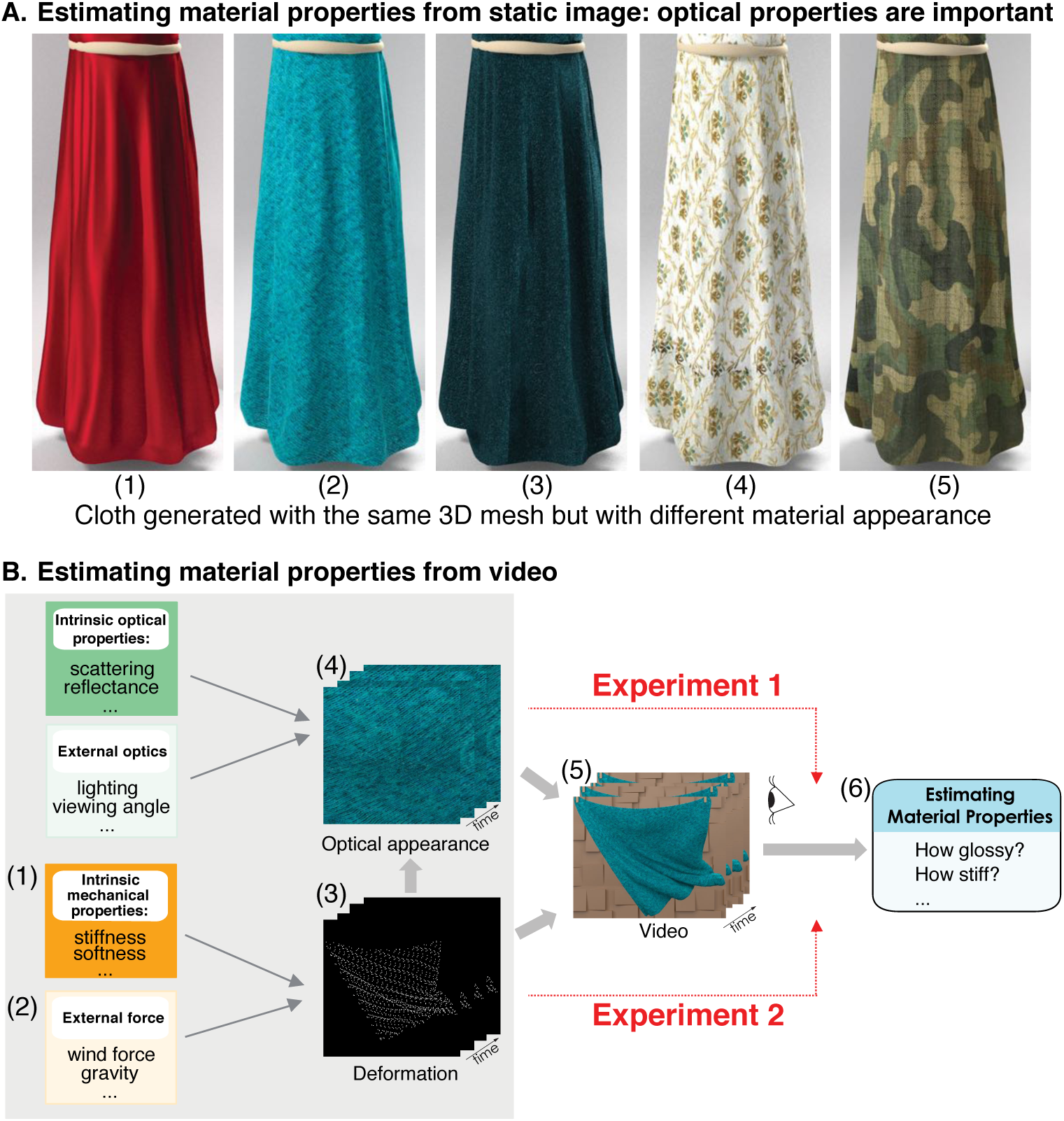
Estimation of the stiffness of cloth from images and videos. A) Cloth samples rendered with the same 3D mesh but with different optical properties. Optical properties affect the perceived stiffness even for clothes with the same intrinsic mechanical properties. B). A generative model for estimating cloth mechanical properties. Both intrinsic material properties and external scene properties influence the optical appearances (4) and deformations (3). In Experiment 1, we aim to compare static image conditions and videos in estimation of stiffness of cloth. Previous studies investigated how stiffness estimation is affected by optical appearance and deformations by directly manipulating intrinsic properties (1) or/and external forces (2). In Experiment 2, we directly manipulate the dynamic deformations (3) by using dynamic dot stimuli and measure how this manipulation affects visual estimation of stiffness.

What information does the visual system use to achieve this? In what aspect does dynamic information help disentangle the contribution of optical properties and mechanical properties? To what degree dynamic deformation alone is sufficient to convey impressions of various mechanical properties? In this paper, we first identified an idiosyncratic deformation pattern (i.e., movement uniformity) to differentiate stiffness, which can be reliably measured by six optical flow features. We further developed a method that directly manipulates the deformation patterns of cloth using dynamic dot stimuli, and showed that manipulating the deformation pattern alone can alter the impression of stiffness of a moving cloth. Using this method, we can examine the causal role of dynamic deformation in stiffness perception. Furthermore, by analyzing optical flow fields extracted from the manipulated dynamic dot stimuli, we confirmed the same six optical flow features can be diagnostic of the degree of stiffness of moving cloth across different scenes.

## Previous work

### Deformation cues in material perception

One of the challenges for understanding material perception is that both the object’s shape and its intrinsic physical properties influence its appearance (Fleming, Jäkel, & Maloney, 2011; Marlow & Anderson, 2015; Marlow, Todorovicć, & Anderson, 2015). This is especially true for deformable objects. In some situations, the visual system can infer material properties just from static shape information without any optical appearances. In a unique study, Pinna and Deiana (2015) showed that simply deforming contours of the object’s boundaries can elicit vivid impressions of different material properties. In our work, we followed a similar reductionism approach by directly manipulating dynamic dot stimuli generated from 3D meshes to alter the perceived stiffness of cloth. Other studies have reported that shape cues are sufficient for the judgment of liquid viscosity and stiffness of deformable and elastic objects (Paulun, Kawabe, Nishida, & Fleming, 2015; Paulun et al., 2017; Spröte & Fleming, 2016). For example, a liquid of a given viscosity can settle into shapes with characteristic features (Paulun et al., 2017; Van Assen & Fleming, 2016; Van Assen et al., 2018) and these features are diagnostic for its viscosity. Previous studies also show that optical properties affect perception of shape deformations(Han & Keyser, 2015; Schmidt, Paulun, Assen, & Fleming, 2017).

Motion has been shown to influence material perception of specular solid objects. For example, Sakano and Ando (2010) showed that the change of the angle of light refraction and reflection caused by head movements is a crucial cue in the perception of glossiness. Doerschner et al. (2011) observed that a moving glossy surface differed from a matte moving surface in three motion features: the coverage, divergence, and 3D shape reliability. They verified the critical role of these motion features by demonstrating that a model trained with these three motion cues can successfully predict both successes and some failures of human material perception. The follow-up study also showed that specular flow is important for estimating 3D shape of glossy objects (Dövenciogğlu, Ben-Shahar, Barla, & Doerschner, 2017). Kawabe, Maruya, and Nishida (2015) analyzed spatialtemporal frequencies of image deformation and have reported that some specific spatiotemporal frequencies allows perception of transparent liquid layers. Most recently, Tamura, Higashi, and Nakauchi (2018) discovered robust dynamic visual cues that could differentiate mirror and glass. Specifically, they found that glass objects had more opposite motion components relative to the direction of the object’s rotation.

Since shape deformation can already tell quite a bit of how deformable an object is under external forces, it is unclear to what degree motion matters. We propose that even though observers can tell a stiff object from soft one from one static frame, this information alone might not be robust over time. When more evidence emerges, observers might change their judgments. As to the perception of elasticity, Kawabe and Nishida (2016) found that human observers were able to recover the elasticity of computer-rendered jellylike cubes based on the shape-contour deformation alone. This was still true even when the cube movies were replaced by dynamic random noise patterns, which retained the optical-flow information but not the surface information. The researchers concluded that the elasticity judgment was based on the pattern of image motion arising from the contour and the optical deformations. In studies regarding liquid viscosity, Kawabe, Maruya, Fleming, and Nishida (2015) extracted the optical flow fields of liquids and presented them as a 2D noise patch array. Even though the shape information has been largely excluded, observers can still distinguish liquids with different viscosity from the noise array. The authors further demonstrated that the visual system utilized image motion speed in the optical-flow field as a cue to estimate liquid viscosity. Additionally, motion is also important for achieving perceptual constancy of mechanical properties in variation of external forces, such as the softness of cloth (Bi & Xiao, 2016) and liquid viscosity (Van Assen et al., 2018).

Recent studies measured the effects of optical properties, shape and motion cues on perception of stiffness and elasticity of deformable objects. Studies in liquid viscosity discovered that viscosity is inferred primarily from shape and motion cues but that optical characteristics influence recognition of specific liquids and inference of other physical properties (Van Assen & Fleming, 2016). Schmidt et al. (2017) investigated the interactions among optical properties, shape and motion cues on the effect of estimation stiffness of unfamiliar objects using material attributes rating task. They found optical appearances elicited a wide range of apparent properties. Using physical simulation, they additionally found that softness of the objects is highly correlated with the extent of deformation. However, when combined the optical cues with the shape deformation, the authors found that optical cues completely dominant. Finally, they presented motion sequences to the observers and found significant effects of motion as well as optical cues. In this paper, we aim to test the hypothesis that dynamic information is highly useful in discounting the effects of optical properties. In addition, different from previous studies, we mathematically manipulate deformation patterns on the dot stimuli generated from the mesh instead of using physical simulation. Our method has the power to dissect which aspect of deformation is critical in stiffness judgment.

### Image synthesis without simulation

Other than computing and identifying image features of perceptual salience, image synthesis is another approach to seek a statistical description of visual appearance that is consistent with human perception. Portilla and Simoncelli (2000) developed a method of synthesizing an image of Gaussian white noise to any given reference texture image. They first built a set of steerable pyramid sub-bands from the reference texture image, and iteratively optimized the sample statistics of each steerable pyramid sub-band and reconstructed an image from the updated pyramid. Similar approach has also been used in synthesizing images of certain material appearance, Kawabe, Maruya, and Nishida (2015) created the impression of transparent liquid by synthesizing the pattern of dynamic image deformation to match the spatiotemporal frequency amplitude spectrum of the resulting pixels to those of the real water. Recently, deep learning has been used to generate images without simulation such as image style transfer (Gatys, Ecker, & Bethge, 2016) and image synthesis of complex materials (e.g., flow) without simulation(Chu & Thuerey, 2017). Particularly, generative adversarial networks (GANs)(Goodfellow et al., 2014) have begun to generate highly photo-realistic images of specific categories, such as faces, album covers, and room interiors (Radford, Metz, & Chintala, 2015), dynamic textures(Xie, Zhu, & Nian Wu, 2017), and videos with scene dynamic (Vondrick, Pirsiavash, & Torralba, 2016). Our method can also be used to modify material properties of stimuli without rendering.

### Cloth perception

Cloth is a common deformable material, yet little is known about how optical properties, shape, and motion affect the perception of its material properties. One debate is how important is dynamic information. Figure 1A shows that optical appearance affects perceived stiffness even for the clothes with the same intrinsic mechanical properties. Most likely observers would perceive the skirt rendered with “silk” optical appearance (Figure 1A(1)) to be more flexible and softer than the one rendered with “velvet” appearance (Figure 1A(3)) even though the two pieces of cloth are rendered with the same 3D models. This might because recognizing the fabric being “silk” biases the perception of it to be less stiff. Yet another explanation might be that the specular highlights on the silk surface could modify the perceived deformation and hence could influence stiffness impression. Aliaga, O’Sullivan, Gutierrez, and Tamstorf (2015) found that the appearance, rather than the motion, dominated the categorical judgment of cloth, except for fabrics with extremely characteristic motion dynamics (i.e., silk). By contrast, other studies find that motion information is important in the perception of mechanical properties of cloth. For example, in a study by Bouman, Xiao, Battaglia, and Freeman (2013), observers estimated the stiffness and mass of cloth examples in real scenes. They found that the observers’ responses were less correlated with the physical parameters in the image condition compared to the video condition. This finding supports the importance of motion in visual estimation of material properties. More recently, Bi, Jin, et al. (2018) reported that when the frame sequences were scrambled, observers’ sensitivity to different stiffness values of cloth was decreased, also suggesting the important role of multi-frame motion information in the perception of stiffness of cloth. To verify this hypothesis, they trained a machine learning model using the dense trajectories, a multi-frame motion descriptor, and demonstrated the robustness of the model in predicting human perceived stiffness of cloth. One reason why we use cloth as model is that its deformation is usually caused by a more complex external force (e.g., oscillating wind) instead of the simple bending or poking force. Inspired by previous studies showing that maximum deformation is a critical cue for stiffness judgment of an elastic cube (Paulun et al., 2017), we conjecture that human visual system might rely on idiosyncratic pattern of dynamic deformation for estimating stiffness, such that under the same external force, a soft cloth deforms to a larger extent but less uniformly than a stiffer cloth.

### Study overview

Figure 1B illustrates the process of how human observers estimate mechanical properties from videos and an overview of our methods. In Experiment 1, we test the hypothesis that humans can partially discount the bias caused by optical appearances and exhibit higher sensitivity to stiffness in video conditions than static image conditions. Second, we analyze the statistics of optical flow fields and discover that cloth with different stiffness values differ in the optical flow movement features relating to the movement uniformity. In Experiment 2, we directly manipulate movement patterns of dynamic dot stimuli generated from the 3D mesh exported from the physics engine. By doing so, we will isolate the dynamic information by removing the influence of optical properties. We investigate whether this manipulation can alter the stiffness judgement and whether this method can be generalized to other scene-setup, 3D model, and generative physics model.

## Experiment 1: Effects of dynamics on stiffness judgment of cloth

Previous research shows that dynamic information is important in judging material properties. It is unclear in what aspect dynamic stimuli have an advantage over static images in the perception of material properties of cloth. We used cloth animations as stimuli and measured the perception of bending stiffness. First, we aimed to show that dynamic stimuli could indeed convey stiffness of cloth better than static images. To do so, we compared stiffness judgments of the same cloth samples displayed in two conditions: static (image) and dynamic (video). We hypothesized that in the static image condition, both intrinsic mechanical properties (i.e., ground truth stiffness value) and surface properties (e.g., textures, surface reflectance) affect stiffness perception. By contrast, in the dynamic condition, the perceived stiffness would be less affected by surface properties and therefore more aligned with the ground-truth values. Second, we investigated what kind of deformation pattern could predict the perceived stiffness by analyzing the optical flow fields. We identified six optical flow motion descriptors that are diagnostic of the degree of stiffness.

## Materials and Methods

### Observers

Eight observers (7 women; mean age = 27.2 years, *SD* = 5.8 years) participated in the experiment on a voluntary basis and were not paid for their participation. All observers reported normal visual acuity and color vision.

### Stimuli

The stimuli were physics-based animations of cloth with various material properties under external forces. Figure 2A illustrates an example of the stimuli in the experiment. On the left is the target cloth and on the right is the reference cloth. In the dynamic condition, both the reference and target cloth were shown as videos; in the image condition, the reference cloth was displayed as videos but the target cloth was displayed as a static image which was a random frame extracted from the corresponding video. The animations were simulated using the Blender Cycles Render Engine (Blender version 2.7.6). The full length of every target and reference cloth video was 8 seconds with 24 frames per second. Each animation differed in one of the following four rendering parameters:

**Figure 2:**
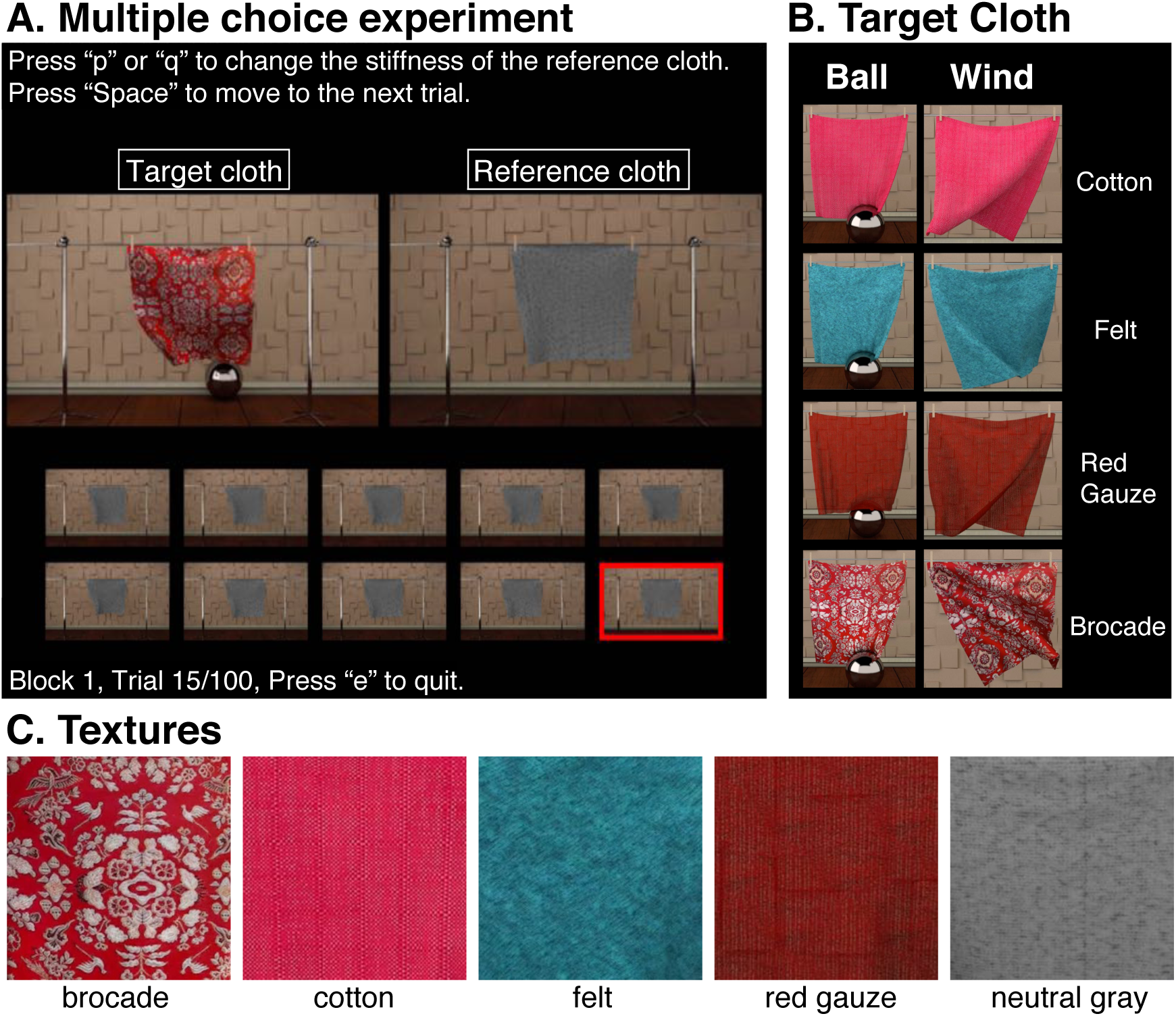
(A) The interface of the multiple choice experiment. Observers adjusted the stiffness of the reference cloth to match that of the target cloth by selecting one of the small reference videos (rows below), which contained the cloth with varying values of bending stiffness. When one of the small reference videos was selected, it would show up at the position of the reference cloth. (B) The target cloth was rendered with four different material appearances (i.e., cotton, felt, red gauze, brocade) and two different scenes (i.e., the ball scene and the wind scene). In the dynamic condition, the target was always presented as video; in the image condition, the target was presented as a single static frame randomly chosen from the corresponding video. (C) A zoom-in view of the material textures. The “neutral gray” refers to the reference cloth, and the rest four belong to the “target cloth”.

### Stiffness

Bending stiffness describes how the cloth forms its wrinkles. Higher values results in bigger but not necessarily more wrinkles. Each target cloth was rendered with one of the five bending stiffness levels {0.01, 0.1, 1, 10, 100}. Each reference cloth was rendered with one of the ten stiffness levels {0.005, 0.01, 0.1, 0.5, 1, 5, 10, 25, 100, 300}, which covered the whole range of stiffness of the target cloth.

### Mass

Mass describes the heaviness of cloth per area. The target cloth had two mass levels {0.1, 0.7}, and each reference cloth was rendered with mass value 0.3.

### Material appearance

Each target cloth was rendered with one of the four distinctive material appearances (see Figure 2B and C): felt, cotton, red gauze, brocade). By contrast, the reference cloth was always rendered with a grayish fleece material (see Figure 2C: neutral gray), which was different from any material appearance of the target cloth. All material appearances were simulated based on the impressions from real cloth samples but not with any particular set of optical parameters, so the resulting videos had different texture, thickness, surface reflectance, roughness, and transparency, making them unique in several dimensions of material appearances.

### Scene set-up

Each target cloth was rendered with one of the two dynamic scenes: a wind scene containing a piece of hanging cloth moving under oscillating wind forces (Figure 2B, right; See Bi, Jin, et al., 2018 for details) and a ball scene containing a rolling ball colliding with a piece of hanging cloth (Figure 2B, left). The reference cloth was rendered with only the wind scene. See supplementary videos for demonstrations of the scenes.

Each observer finished 160 trials in both the static and dynamic conditions (5 stiffness levels × 2 mass levels × 4 material categories × 2 scene set-ups × 2 repetitions).

### Procedure

Observers were presented with videos and images of cloth simulations and compared their judgments of stiffness. Figure 2A shows the interface of the multiple choice task used in this experiment. During each trial, the observers adjusted the stiffness of a reference cloth by selecting one of the ten small reference videos below to match the stiffness of a target cloth. The ten small reference videos were rendered with varying bending stiffness values, which covered the whole range of the stiffness from the least stiff to the most stiff. When one of the small reference videos was selected, it would show up as a full-sized video at the position of the reference cloth and replaced the default video. At each trial, the target cloth varied in one of the following four factors: stiffness, mass, the material appearance, and scene set-up. We used the same set of small reference videos for each target video across all trials.

## Results

Figure 3 plots the mean (A and B) and median (C and D) matched stiffness across all observers versus the ground-truth stiffness values of the target cloth. First, we found that in both image and video conditions, observers could distinguish cloth with different bending stiffness values. Across all conditions, the matched stiffness increases as the ground-truth stiffness of the target cloth increases. Second, the figure shows that the lines in the video condition are steeper than those in the image condition, which is especially obvious when plotting the median (Figure 3 C and D). This suggests that observers showed higher sensitivity to different stiffness values in the dynamic condition than in the static condition.

**Figure 3:**
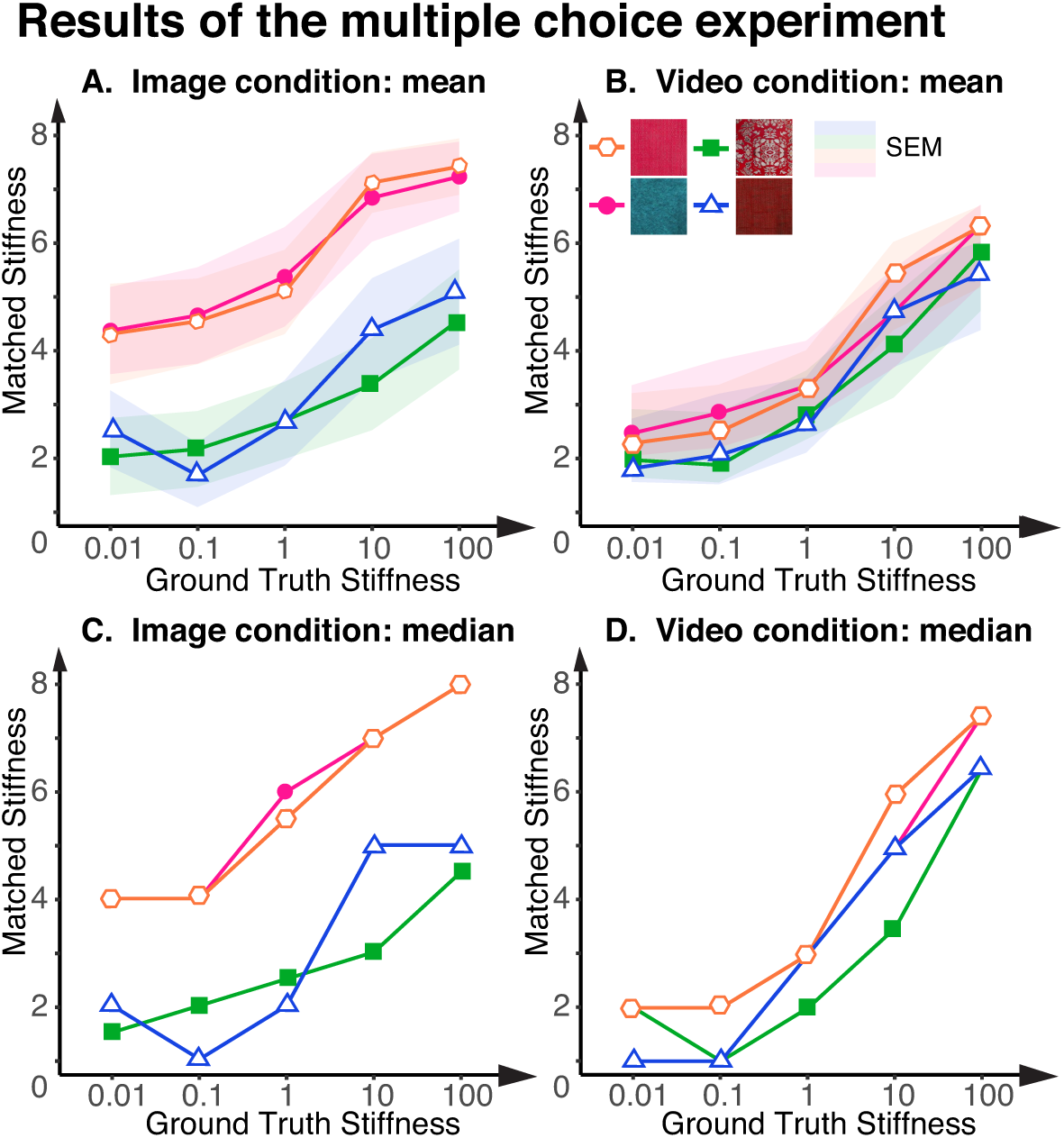
Matched stiffness plotted as function of ground-truth stiffness levels in image and video conditions. The x-axis shows the ground-truth stiffness value of the target cloth. The y-axis shows the matched stiffness levels. Different colors indicate different material appearances. (A) Mean matched stiffness levels plotted as function of ground-truth stiffness for image condition. (B) Mean matched stiffness levels plotted as function of ground-truth stiffness for video condition. (C and D) Same as A and B, but the matched stiffness levels are plotted using the median value across all observers.

We analyzed observers’ performance in the experiment by a two-way repeated-measure analysis of variance (ANOVA) on the matched stiffness with the material appearances (four levels) and the stiffness values (five levels) of the target cloth as fixed within-subject factors. We did separate analysis for the video and image conditions.

The main effect of *material appearances* was signifiant in both the image condition (*F*(3,21) = 14.73, *p* < .0001) and the video condition (*F*(3,21) = 3.38, *p* = .038). Post-hoc analysis indicated that cotton and felt were perceived to be stiffer than red gauze and brocade(*p*s < .05). But the effect largely decreased in the video condition (*η*^2^ = 0.026) when compared with the image condition (*η*^2^ = 0.366).

Additionally, the main effect of *ground-truth stiffness of the target cloth* on the matched stiffness was significant in both the image condition (*F*(4,28) = 33.69, *p* < .0001) and the video condition (*F*(4,28) = 41.94, *p* < .0001). But the effect is much larger in the video condition (*η*^2^ = 0.624) than in the image condition (*η*^2^ = 0.287).

### Optical flow analysis

The perceptual results show that in the video condition, observers show enhanced sensitivity to stiffness, and furthermore, the effect of material appearances has been partially discounted. This suggests that the videos must contain information that allows observers to estimate the intrinsic stiffness without being affected much by material appearances. To determine what kind of motion information is available and important for the inference of stiffness, we analyzed the statistics that extracted from the optical flow fields of the cloth videos, using the method described in Kawabe, Maruya, Fleming, and Nishida (2015). Some of these statistics have been shown to be highly correlated with the perceived liquid viscosity and the stiffness of cloth (Bi & Xiao, 2016; Kawabe, Maruya, Fleming, & Nishida, 2015).

Figure 4A shows that the optical flow fields of a soft moving cloth is different from that of a stiff one. The two cloth videos were rendered with all other parameters exactly the same except the bending stiffness values. We observed that the flow vectors of a soft cloth are less uniform in both the direction and the magnitude than those of a stiff cloth. This motivated us to look for motion features that describe such movement uniformity. To do so, we compared the optical flow statistics between a soft cloth (bs = 0.1; solid lines in Figure 4B) and a stiff one (bs = 100; dotted lines in Figure 4B), with the wind forces in all cloth videos kept the same. We analyzed the 16 motion descriptors that were proposed by Kawabe, Maruya, Fleming, and Nishida (2015) and found that the mean and standard deviation (SD) of three motion features (Divergence, Gradient, and discrete Laplacian) could typically differentiate the stiffness values and might account for the movement uniformity. Specifically, Figure 4B shows that the videos containing a less stiff cloth typically shows higher values in all these statistics than the videos containing a more stiff cloth, regardless of optical appearances. The detailed calculation of these statistics can be found in Kawabe, Maruya, Fleming, and Nishida (2015).

**Figure 4:**
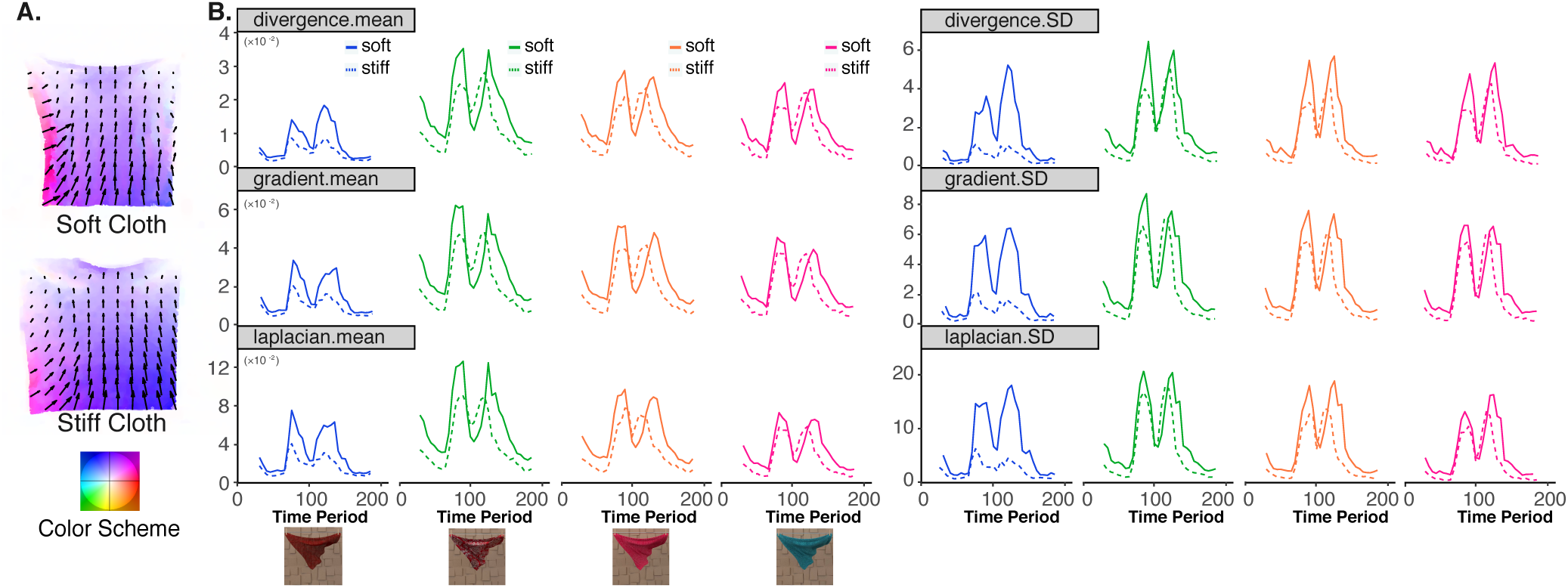
Optical flow analysis of videos containing simulated cloth. (A) Two-frame optical flow vectors are less uniform for a soft cloth (upper panel) than a stiff cloth (lower panel). The color maps are plotted together with the displacement vectors to show the length and direction of the displacements. In the vector fields, larger vector refers to more displacement and the arrow points to the movement direction. In the color map, the saturation indicates the magnitude of the movement, while the hue represents the direction of the movement. (B) Comparison of optical flow statistics between a soft cloth (bs = 0.1; solid line) and a stiff one (bs = 100; dotted line). Different colors indicate different material appearances. Across the whole time period that has been plotted and for all plots, the solid lines are typically above the dotted lines, indicating that all the optical flow statistics have higher values for the softer cloth compared to the stiffer one.

The optical flow analysis shows that the idiosyncratic deformation pattern (i.e., movement uniformity) measured by the six optical flow features can be used to differentiate stiffness of cloth in variation of optical appearance. However, the extraction of optical flow field can be easily affected by optical properties. In order to directly test how the pattern of dynamic deformation affects the stiffness judgement, we propose to isolate the deformation pattern by removing the influence of optical properties. Hence, in the next section, we showed a method to directly alter the movement pattern by manipulating the displacement vectors in the 3D mesh of the cloth, and demonstrated that changing the pattern of dynamic deformation alone can alter the perceived stiffness.

## Experiment 2: Patterns of dynamic deformation affect inferred stiffness

In Experiment 1, we discovered that motion statistics associated with movement uniformity could be diagnostic of degree of stiffness. This finding motivates us to propose a method to isolate and manipulate the dynamic deformation that is related to movement uniformity, and to demonstrate that this manipulation alone can alter the inferred stiffness.

To isolate the dynamic deformation information, we created dynamic dots from the 3D mesh of a moving cloth as stimuli (Figure 5). In the dynamic dot stimuli, the movements are generated by the displacement of each dot. We can manipulate the pattern of dynamic deformation by directly varying the displacement vectors. One way to systematically vary the displacement vectors is to vary the moving velocity. We name the uniformity of the velocity of the dots as “velocity coherence”, which describes the movement uniformity.

**Figure 5:**
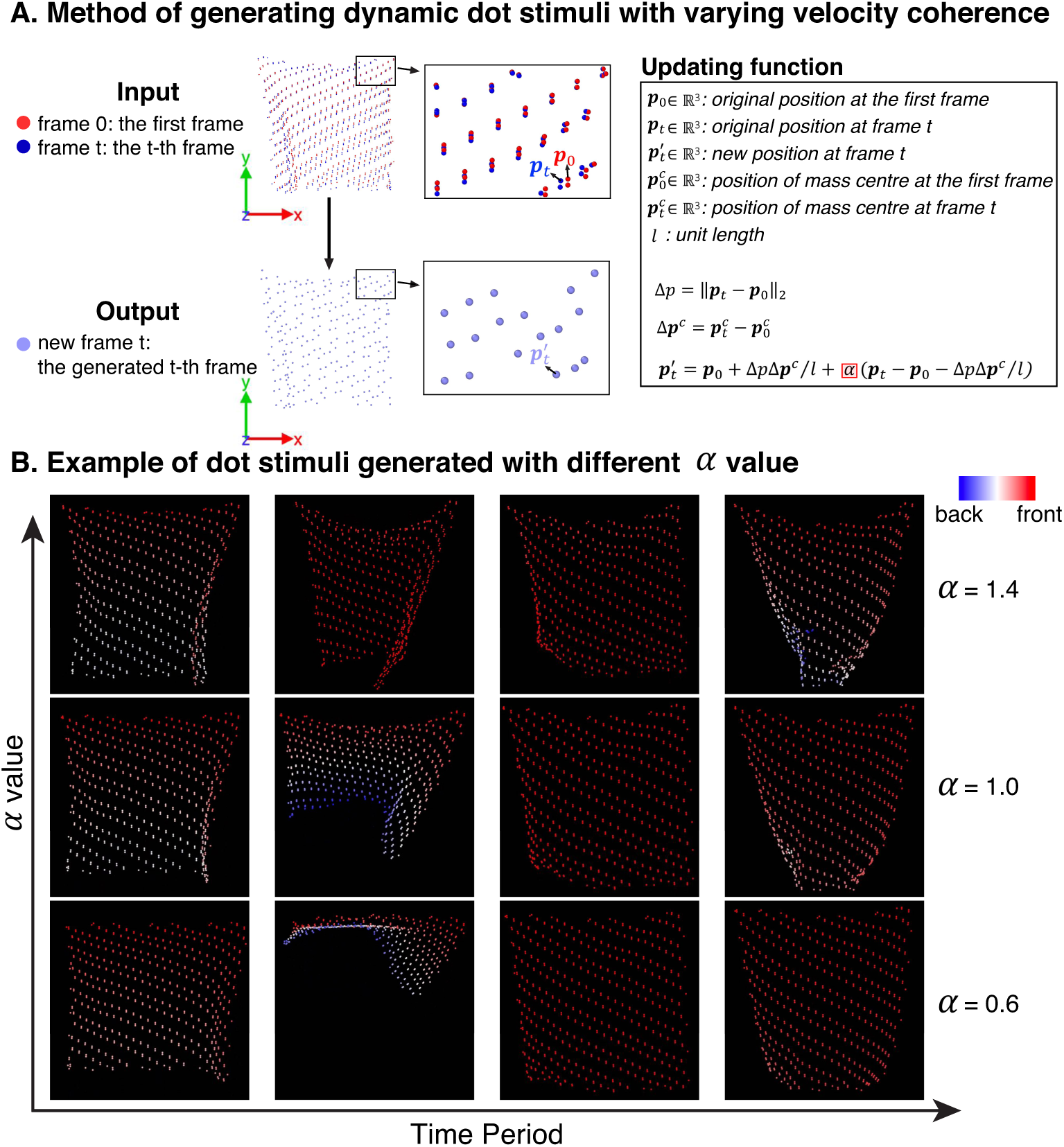
Experiment 2: Using dynamic dot stimuli to isolate and manipulate the patterns of dynamic deformation. (A) Method of creating the dynamic dot stimuli. The input frames are exported from Blender. The output frame is generated by shifting the positions of the dot in the original frame by the updating function defined on the right box. For each dot in the original frame *t*, its new position 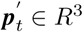 in the new frame *t* is updated using both its current position (***p*** *∈* R^3^) as well as its position at the first frame (***p*** ∈ *R*^3^) (right panel). The *α* in the function determines the velocity coherence. Smaller *α* value makes the dots to move more uniformly. (B) Examples of the dot stimuli generated with different *α* values. X-axis represents the time period. Y-axis represents three *α* levels. The color hue represents the depth information.

The velocity is defined as the spatial displacement of one dot within every two consecutive frames. The dynamic dots move more uniformly when a large number of dots move with similar velocities. In contrast, the movement uniformity is low when the velocity of the dots differs a lot among each other. In Experiment 2, we aim to test the hypothesis that movement uniformity affects the perceived stiffness of dynamic dotted cloth videos, such that increase the movement uniformity makes the dotted cloth appear stiffer.

## Materials and Methods

### Observers

Eight observers (6 women; mean age = 24.75 years, *SD* = 4.3 years) participated in the study on a voluntary basis and were not paid for their participation. Four of them also participated in Experiment 1.

### Stimuli

Figure 5 shows examples of dynamic dot stimuli and the method we used to generate them. To generate the dot stimuli with different movement uniformity, we first need a template dot video, which was the 3D mesh outputs from Blender animations of a moving cloth in the wind scene as described in Experiment1 (see Figure 2B, wind scene). The outputs were 200 sequential wavefront .obj files, and by default, the cloth 3D mesh in each frame contained 21,626 vertexes (i.e., dots). We then reduced the number of vertexes in each frame to 664 using the systematic sampling method to make the dot stimuli appear less dense but still appeared like a piece of cloth.

Based on the template video, we generated new dynamic dot stimuli with different movement uniformity by directly manipulating the displacement of each dot, using the updating function shown in Figure 5A. Specifically, for each dot in any given frame *t*, its new position 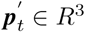 in the new frame *t* was calculated using both its current position (***p***_*t*_ ∈ *R*^3^) as well as its position in the first frame (***p***_0_ *∈ R*^3^). The *α* in the function determined different movement uniformity. When *α* < 1, the updating function decreases the displacement amplitudes of dots that move more than average, and increase the displacement amplitudes of dots that move less than average, therefore making the movement more uniform. On the other hand, when alpha *α* 1, the updating function increases the displacement amplitudes of dots that move more than average, and decrease the displacement amplitudes of dots that move less than average, leading to decreased movement uniformity. Overall, smaller *α* value corresponded to increased movement uniformity, which would make the dot stimuli appear stiffer, as we hypothesized.

To experimentally test this hypothesis, we generated eight dynamic dot videos with uniformly sampled *α* values {0.2, 0.4, 0.6, 0.8, 1.0, 1.2, 1.4, 1.6} and conducted a psychophysics experiment to measure the perceived stiffness of the object represented by these dots stimuli. If the perceived stiffness is correlated with the *alpha* values in the dot stimuli, we then have the evidence to show that movement uniformity was useful in estimating stiffness.

### Design and Procedure

We used Maximum Likelihood Difference Scaling (MLDS) with the method of triads (Maloney & Yang, 2003; Knoblauch, Maloney, et al., 2008) to measure the psychometric function relating changes in the *α* value to changes in perceived stiffness by humans. In each trial, observers were presented with a triplet of videos and asked to judge which video pair (left and center) versus (right and center) appear more *different* from each other in terms of stiffness. They indicated their choice by pressing the P (left pair) or the Q key (right pair). On any given trial, the three videos in the triads always had different *α* values, and the *α* values of the center videos were always between those of the left and right ones. Therefore, the movement uniformity of the three videos was either in ascending (left < center < right) or descending (left > center > right) order.

## Perceptual results

The perceptual scale for each observer was computed using the MLDS package for R from Knoblauch et al., 2008. Figure 6A shows the estimated perceptual scale for each observer as a function of the *α* value, along with the mean across all observers, which were estimated by MLDS using the GLM (generalized linear model) implementation (McCullagh, 1984). There was a strongly negative correlation between the perceptual scale and the *α* value (*r*(64) = −0.97), indicating that observers were able to distinguish the dot stimuli with different velocity coherence values. A linear regression fitted to these data revealed that the perceived stiffness decreased as the *α* increased in a significant linear fashion (F(1,62) = 1060.7, *p* < .0001). Together, in support of our hypothesis, higher movement uniformity made the cloth in dynamic dot stimuli appear stiffer.

**Figure 6:**
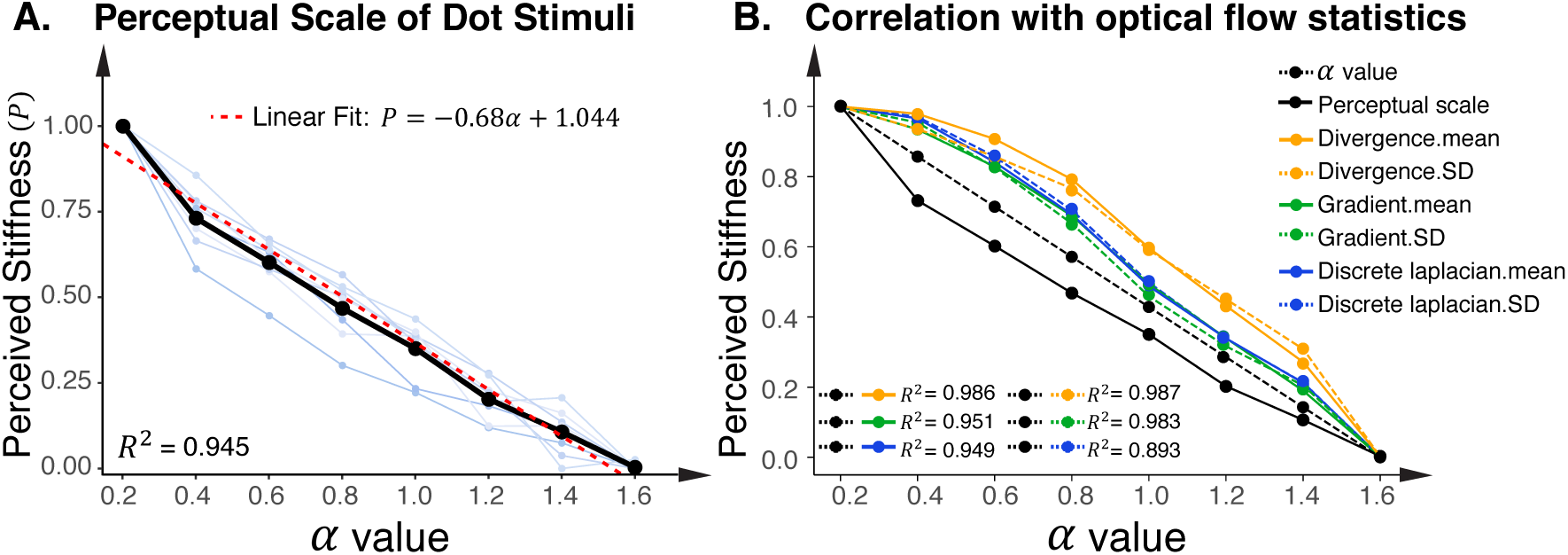
Results of Experiment 2: (A) Measured perceptual scale of stiffness from the dot video as function of different *α* values. The black line represents the scale averaged over the 8 observers, and the blue lines represent individual observer’s scales. X-axis represents *α* values of different dynamic dot stimuli. Y-axis represents the averaged perceptual scale of stiffness. Perceived stiffness decreases as the *α* value increases in a linear fashion. The dotted red line refers to the linear fit between the perceptual scale and the *α* value.(B) Optical flow statistics and perceptual scale are plotted as function of *α* values. X-axis represents *α* values of different dynamic dot stimuli. Y-axis represents the normalized values of the perceptual scale as well as the six optical flow statistics averaged across time. The *α* value, optical flow statistics, and perceived stiffness from the dot stimuli are highly correlated with each other.

### Optical flow analysis

We have demonstrated that directly manipulating the pattern of dynamic deformation can alter the perceived stiffness of cloth in the dynamic dot stimuli. Specifically, the cloth in dot videos is perceived to be stiffer when the dots move more uniformly. Next, we would like to confirm that the same six optical flow features that proposed in Experiment 1 could also be disgnostic of the degree of stiffness of dot stimuli.

To verify this, we used the same method as Experiment 1 to extract the optical flow statistics from the dynamic dot stimuli. Figure 7 plots the mean and standard deviation of divergence, gradient, and discrete laplacian of the eight dynamic dotted stimuli that were generated with different *α* values. The figure shows that the blue lines are typically above the yellow lines, suggesting that the values of optical flow features increases as the *α* value increases (i.e., perceived to be softer). This is consistent with the results of the optical flow analysis in Experiment 1.

**Figure 7:**
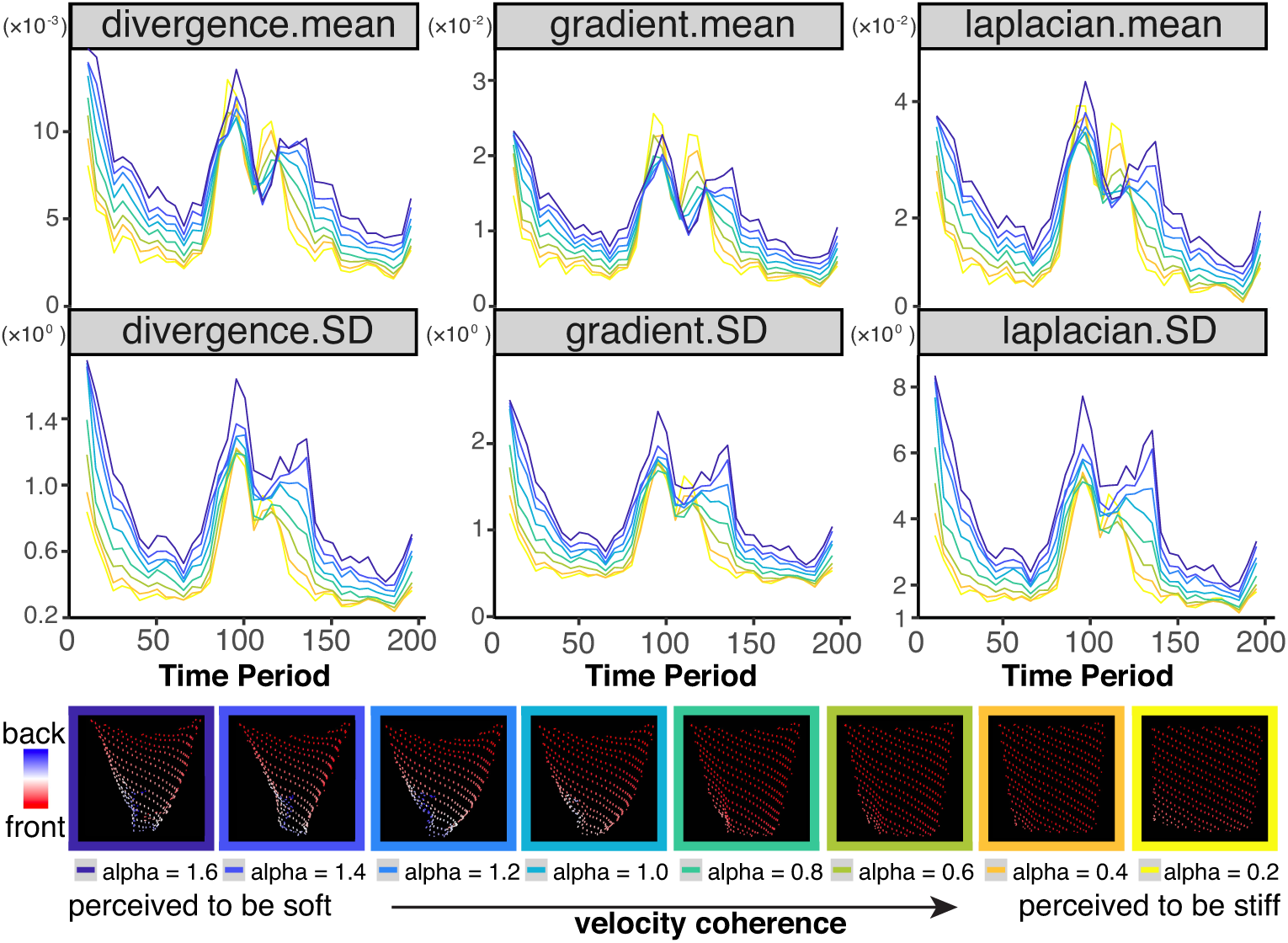
The optical flow statistics extracted from the eight dynamic dot stimuli with different *α* values. Each line represents optical flow statistics of a specific *α* value. Across the majority of the time period that has been plotted (i.e., 0 *∼* 200 frames), the bluish lines are above the yellowish lines, indicating that all the six optical flow statistics are higher for the dot stimuli with higher *α* values (i.e., perceived to be softer).

Further correlation analysis demonstrated that the six optical flow statistics were highly correlated with both the perceived stiffness and the *α* value (*r*^2^s > 0.89; see Figure 6B). Together, these findings supported our hypothesis that when judging stiffness of cloth in videos, human observers might rely on the idiosyncratic pattern of dynamic deformation such that a soft cloth moves less uniformly than a stiff cloth. More importantly, such pattern of dynamic deformation can be directly manipulated by the updating function we proposed, and can be measured by the optical flow features extracted from the videos.

### Test of robustness

It is possible that our finding is particularly restricted to the cloth in the wind scene, because the movement pattern as well as the optical flow fields would be highly affected by the scene set-ups. One can also argue that our finding is limited to the generative physics model used to simulate the cloth. Therefore, we used a different physics model to simulate cloth in two different dynamic scenes. In addition, we aim to test whether this method could also alter the perceived softness of a different 3D object such as an elastic cube.

#### New generative physics model and scene set-up

To account for these, we used Baraff and Witkin (1998) method to model cloth dynamics, which is different from the cloth model used by Blender (Provot et al., 1995; Provot, 1997). Specifically, based on the implementation by Pritchard (Freecloth 0.7.1), we exported the 3D dot cloth animation sequences under two new dynamic scenes: the drape scene contained a piece of cloth draping over a square desk (Figure 8A, “3D mesh” panel), and the corner scene contained a piece of cloth hanging over, with three corners pinned and the other corner being released (Figure 8B, “3D mesh” panel). We used the same method as described in the stimuli section in Experiment 2 to create the dynamic dot stimuli with different *α* values. As Figure 8 (A and B) shows, the dot stimuli with a higher *α* value appeared softer than those with a lower *α* value. Additionally, consistent with the main findings in Experiment 1 and 2, the dynamic dot stimuli with lower *α* value had smaller values in all of the six optical flow statistics that we proposed.

**Figure 8:**
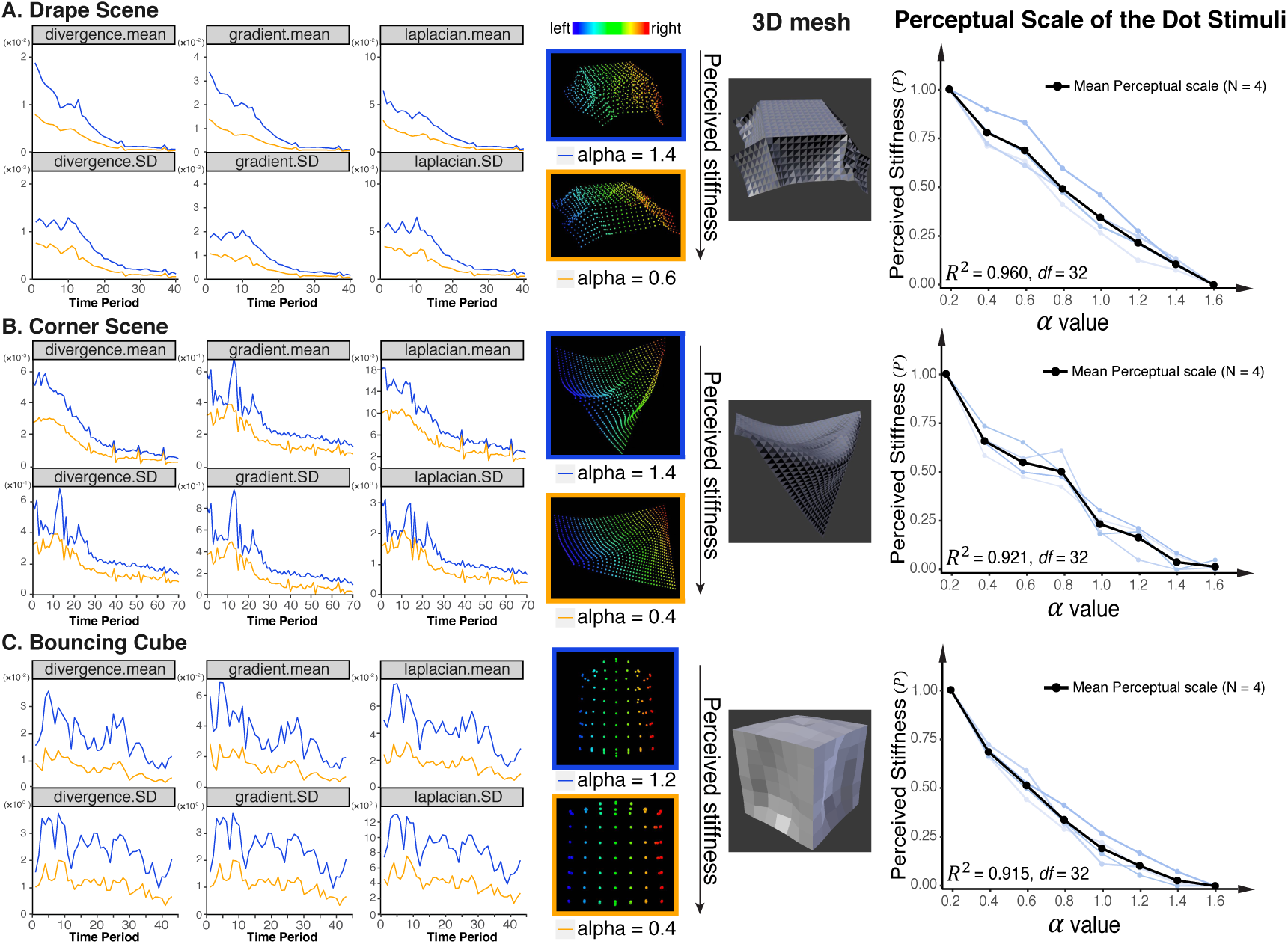
Results of optical flow analysis of dynamic dot stimuli in three new conditions: cloth in the drape scene (A) and in the corner scene (B), as well as an elastic bouncing cube (C). Each line in the leftmost panel represents the optical flow statistics value as function of time. Blue colored line represents high *α* values while orange colored lines represent low *α* values. For all of these three conditions, the blue lines are above the orange lines. This indicates that the six optical flow statistics are higher for the dot stimuli with higher *α* values. The rightmost panel plots the measured perceptual scales from four new observers. Similar to the findings in Experiment 2, the perceived stiffness is highly correlated with the *α* values.

To measure the perceptual scale of the dot stimuli in the two new scenes, we sampled 8 *α* values (same as Experiment 2) and conducted a MLDS experiment on four new observers. In both scenes, the measured perceptual scales of stiffness (Figure 8A and Figure 8B: rightmost panel) were highly correlated with the *α* values (*r*(32)^2^ > 0.92).

#### New 3D model

Next we were interested to know whether our findings could be generalized to other non-rigid objects. To answer this, we created an elastic cube bouncing on the ground in Blender (see Figure 8C, “3D mesh” panel). Next we used the same method to modify the pattern of dynamic deformation and generated the dynamic dot stimuli. We then extracted optical flow statistics. Figure 8C shows that the results are consistent with previous ones: the dynamic dot stimuli with higher velocity coherence had smaller values in all of the six optical flow statistics. Similarly, results of an MLDS experiment reveals that the measured perceptual scale of the dot stimuli was highly correlated with the *α* value (*r*(32)^2^ = 0.915).

Together, these results suggest that our findings are can be generalized to other dynamic scene set-up, generative physics model, and other deformable object.

## Discussion

This article aimed to understand whether and how dynamic deformation affects the inference of stiffness of deformable objects in dynamic scenes. First, we found stiffness perception was highly affected by optical appearances when the cloth was viewed as static images. However, when presented as videos, the effect of optical appearances had been partially discounted, leading to enhanced sensitivity to different stiffness values. Optical flow analysis showed that motion statistics associated with movement uniformity (i.e., divergence, gradient, and discrete laplacian) could be diagnostic of the degree of stiffness when the appearances were the same. In Experiment 2, we demonstrated that directly manipulating patterns of the dynamic deformation of a dot cloth video using a mathematical function alone could alter the perceived stiffness. We could use the same method to alter the perceived stiffness of deformable objects across a variety of scene set-ups, 3D model, and generative physics model. Finally, we confirmed the same six low-level optical flow image features could discriminate the degree of stiffness from the dot stimuli.

### Estimating mechanical properties using image-based features

Intrinsic mechanical properties interact with external forces in a complex way to affect the movement of cloth in the video. Due to the complexity of this generating process, it is unlikely that human can inverse this process to accurately estimate the mechanical properties. More likely, human visual system can infer stiffness by identifying image-based features (i.e., features that directly extracted from images/videos) that are more affected by intrinsic stiffness than other factors.

Using the framework of the *statistical appearance model*, one line of work investigated the role of verbally reported mid-level features. For example, Schmidt et al. (2017) reported that perceived deformation is important in stiffness perception of unfamiliar objects. Van Assen et al. (2018) found the perceived liquid viscosity could be well predicted by four factors, which were verbal judgments relating to the distribution, irregularity, rectilinearity, and dynamics. Even though these studies help understand the perception of mechanical properties, they didn’t identify what image-based measurements that the visual system might use to represent these verbally reported features.

Another line of studies have identified multiple image-based cues in perception of deformable materials that might be used by human observers. Kawabe, Maruya, Fleming, and Nishida (2015) computed optical flow fields between two consecutive frames of running transparent liquid and decomposed into horizontal and vertical vector components. After Fourier transform of each of the decomposed components, they find that the amplitudes of spectra of image deformation, but not the phase, is critical for identifying a transparent liquid. They also identified critical spatial temporal frequencies that are correlated with impression of transparent liquid. Studies by the same author also showed that motion speed is critical for perception of liquid viscosity (Kawabe, Maruya, Fleming, & Nishida, 2015) and judgement of elasticity deformable transparent cubes (Kawabe & Nishida, 2016). Previous work showed that image motion speed can also be used to discriminate the rigid from non-rigid cylinders (Jain & Zaidi, 2011). Nevertheless, other studies find that mean speed is not enough. For example, it was found that the magnitude of phase differences in the oscillating motion could influence the visual impression of an illusory object’s elasticity (Masuda et al., 2013; Masuda, Matsubara, Utsumi, & Wada, 2015).

Here, we discuss the role of motion speed in perception of stiffness from the dynamic dot stimuli used in our study. In Figure 9A, we plotted mean speed of optical flow vectors for dot stimuli with different alpha values. The figure shows that the image motion speed alone is insufficient to discriminate the stiffness during slow variation of external forces (see red circled area in Figure 9A). Compared to other six optical flow statistics that we used in this paper (Figure 9E), the image motion speed can better discriminate the stiffness during rapid variation of the external forces but is worse in discriminating the stiffness when the external forces are slowly-varying. Our results suggest motion speed and features associated with movement uniformity might be supplementary to each other in discriminating stiffness.

**Figure 9:**
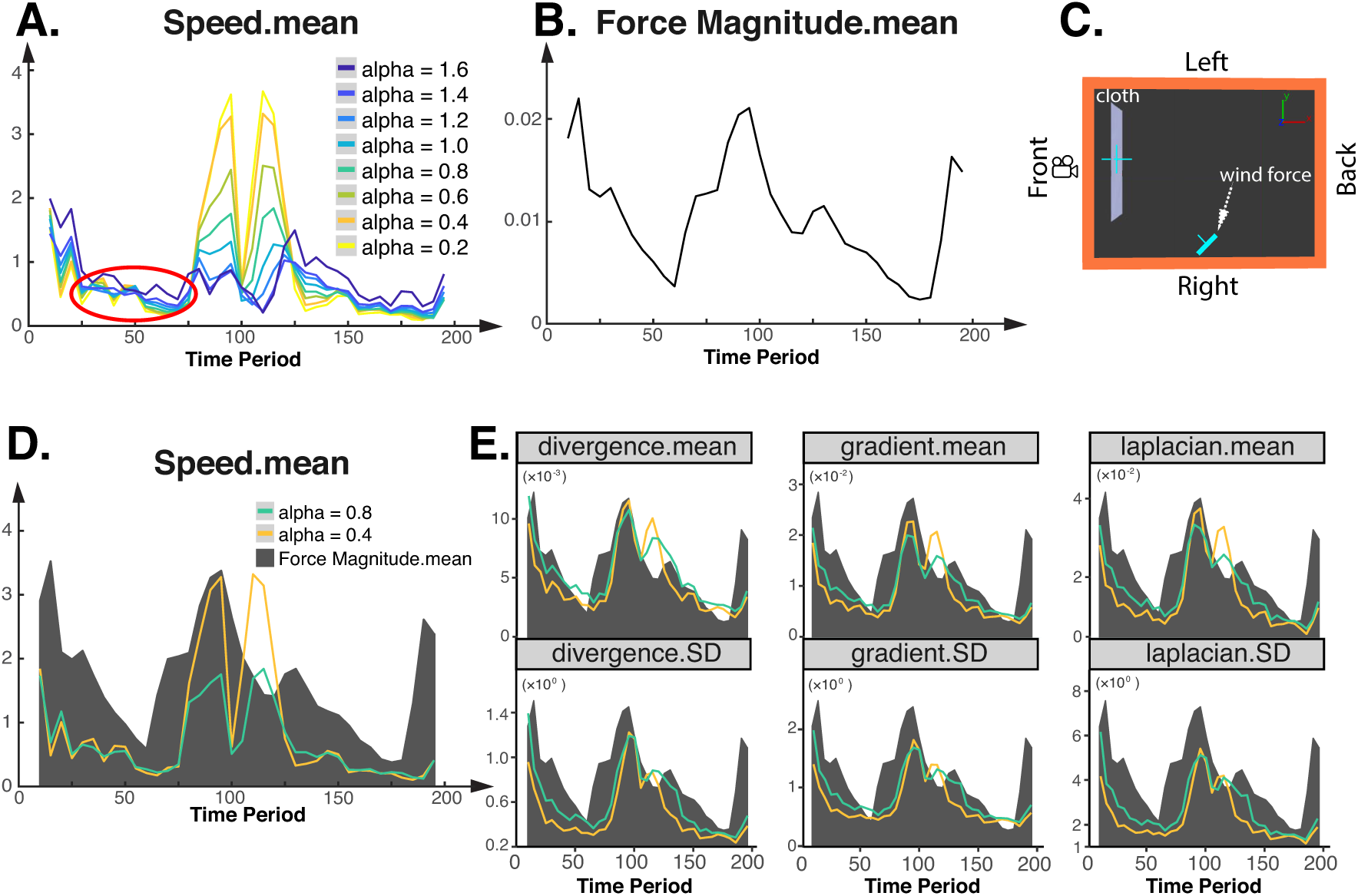
Comparing mean speed (mean norms of optical flow vectors) and other optical flow statistics in stiffness discrimination across the time period of applied force. A. Mean speed of the optical flow vectors for dot stimuli with different *α* values. The area in the red circle is an example period during which the mean speed is not sufficient in distinguishing different stiffness levels. B. The mean magnitude of received force of the cloth across the time period. C. Configuration of the scene geometry. The location of the wind is fixed but it rotates around the cloth periodically. D. Example plot of the mean speed of optical flow vectors for dot stimuli of two *α* values along with the magnitude of received force across the time period. The mean speed can well discriminate the stiffness during the rapid variation of the force (e.g., 75th frame to 125th frame) but cannot reliably discriminate the stiffness when the force varies slowly (e.g., 25th frame to 75th frame). E. Other optical flow statistics for dot stimuli of the same alpha values and the magnitude of received force across the time period. These statistics can discriminate the stiffness better during the slow variation of external forces.

### Static deformation versus dynamic deformation

Previous works show that the stiffness perception of deformable objects is dominantly affected by the absolute magnitude of deformation and translation (Warren Jr, Kim, & Husney, 1987; Paulun et al., 2017; Schmidt et al., 2017). For example, in a dynamic scene that contains a bouncing ball, the perceived elasticity of the ball is barely affected by the velocity information, but mainly determined by the absolute translation information such as relative height (Warren Jr et al., 1987). More recently, Paulun et al. (2017) used computer-rendered animations of an elastic cube being pushed downward to various extents and demonstrated that perceived elasticity was mainly determined by the absolute magnitude of deformation compared to the original non-deformed state, regardless of whether the stimuli were presented as static images or videos. They concluded that, to judge stiffness, the human visual system might rely on the extent to which an object changes its shape under external forces.

The update function that we proposed in this study changes both the dynamic deformation (i.e., movement uniformity) as well as the maximum deformation (i.e., the maximum magnitude of deformation). It could be argued that the maximum deformation alone is enough for the inference of stiffness and the pattern of dynamic deformation does not provide more information. We conjecture that the maximum deformation will dominantly affect the perceived stiffness when the two pieces of fabrics are very different in their maximum deformations. Humans tend to judge the fabric with larger maximum deformation to be softer than the one with smaller maximum deformation. However, when the two pieces of fabrics’ maximum deformations are similar, humans might rely on the pattern of dynamic deformation to distinguish stiffness.

With a small modification to the updating function, we created a demo to provide preliminary support for the importance of the pattern of dynamic deformation in inferring stiffness. Specifically, we partially isolated the maximum deformation and the movement uniformity, and created two new cloth videos (see supplementary materials): one moves more uniformly but with larger deforming speed, and the other one moves less uniformly but with smaller deforming speed. If stiffness judgment only relies on the absolute magnitude of deformation, observers would always judge the cloth with larger speed to be softer. However, as reported by three new observers, the one that moves less uniformly was reported to be softer, indicating that the effect of movement uniformity outweighed that of the absolute magnitude of deformation. Interesting, the observers reported the cloth video with larger deforming speed to be blown by stronger winds. This pilot data suggests that in a complex scene that has varying and unknown forces, the stiffness perception of cloth might rely on both the static deformation (i.e., absolute magnitude of deformation) as well as the pattern of dynamic deformation.

### Implications in neuroscience, computer vision and graphics

Our method can be potentially applied to other areas. Since the dynamic dot stimuli are spatially simple and the dynamic deformation pattern is quantifiable, this could potentially serve as a paradigm to study the role of visual motion on material perception of objects in neuroscience. In addition to the optical flow features that discussed in this paper, there are many other image features that can represent visual motion. To see whether other physiologically plausible computations based on spatiotemporal filters can be used to account for our results, we computed outputs of linear combination of motion energy features from the dynamic dot stimuli (Adelson & Bergen, 1985; Nishimoto et al., 2011; Vinken, Bergh, Vermaercke, & Beeck, 2016). Figure 10A illustrates the details of the model. First, we convolved the video frames with a bank of quadrature pairs of Gabor filters, each with a certain spatiotemporal frequency and orientation. The output of each quadrature pairs was then squared and summed to give the energy features. Figure 10B shows that the simplest linear combination (i.e., equal weights) can well discriminate dotted cloth videos with different alpha values. Future studies can examine the neural responses when these dotted videos are presented to the visual system and determine the neural basis of discriminating stiffness from videos.

**Figure 10:**
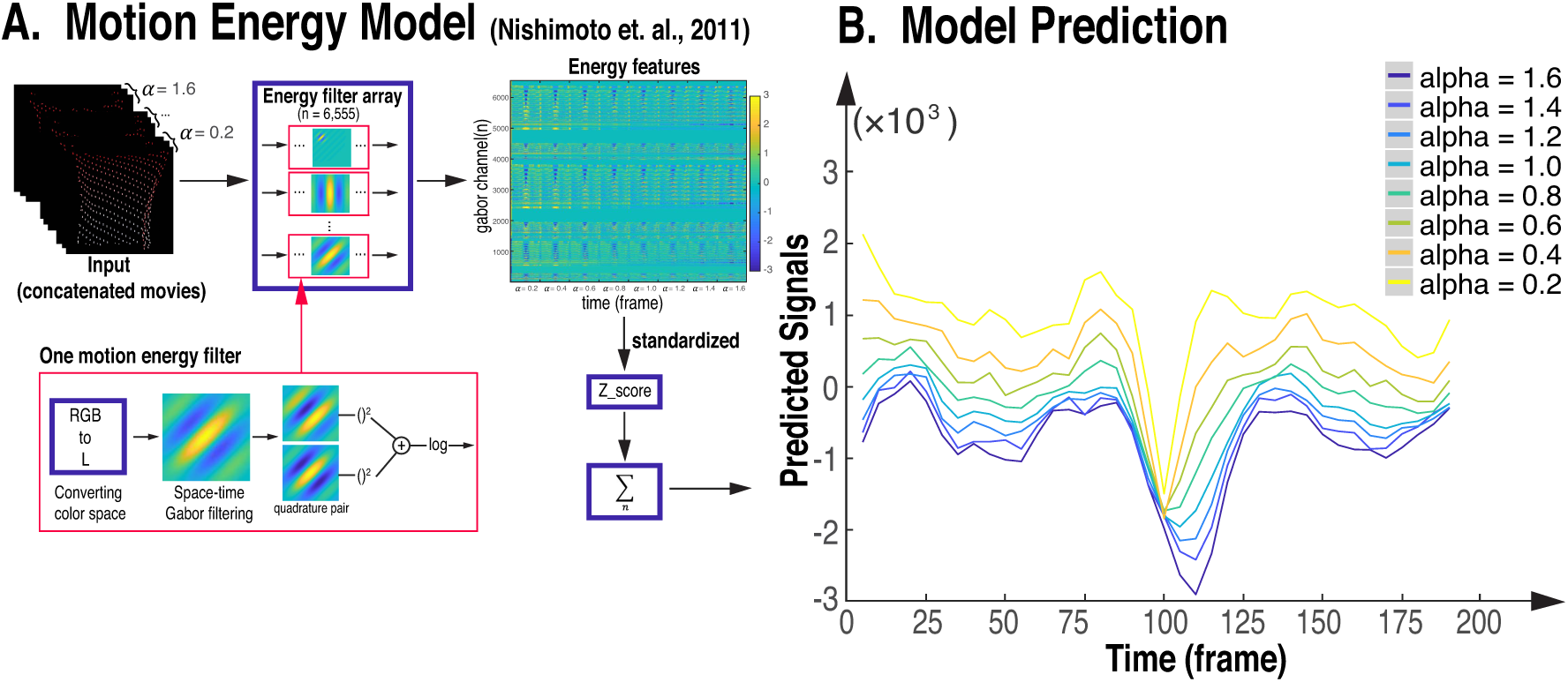
Linear combination of the motion energy features can also be used to discriminate stiffness from the dot stimuli. (A). Flow chart of the motion energy model. Cloth videos with different alpha values were concatenated as input stimuli. We first converted the input to LAB color space and then convolved the luminance channel of the input stimuli with a bank of quadrature pairs of Gabor filters, each with a certain spatiotemporal frequency and orientation. The output of each quadrature pair is then squared and summed to give energy features. And the output from all Gabor filter channels is standardized, summed up and plotted as predictions in B. (B). Outputs from the linear combination of motion energy features with eaqual weights for the dot stimuli with different *α* values across the time period. The prediction by motion energy features can also be diagnostic of degree of stiffness.

Previous work suggests that visual motion often provides powerful cues for scene segmentation for both human and artificial systems (Wagemans et al., 2012; Shi & Malik, 1998). More recently, an fMRI study finds that when showing a class of bistable moving stimuli, the perceptual scene-segmentation was associated with an increased activity in the posterior parietal cortex (PPC) together with a decreased neural signal in the early visual cortex (Grassi, Zaretskaya, & Bartels, 2018). This suggests that PPC is a hub involved in structuring visual scenes based on different motion cues. Using our dynamic dot stimuli, we could test the hypothesis that PPC is involved in estimating stiffness from movement patterns. This will further advance the search for neural substrate for material perception. From the computer vision perspective, our method could serve as a fast way to modify the stiffness of non-rigid objects, which can benefit real-time rendering and apparent motion modification. Most recently, Punpongsanon, Iwai, and Sato (2018) used a similar method to efficiently change the perceived fabric bending stiffness using an optical flow enhancement technique in augmented reality. Specifically, they first extracted apparent motion from a real cloth, then directly modified the apparent motion. Using a projector to map the new apparent motion back to the original fabric, they were able to change the perceived stiffness of the real fabric. Future studies could modify our method into a standard way to change the perceived stiffness of general non-rigid objects in a wide range of scenes. The stiffness parameter used in most physical engines (e.g., Blender) is extremely non-linear. Thus, in the previous studies, the authors have to sample the parameters based on their experiences to generate stimuli with varying stiffness levels.

Our work could also contribute to computer graphics by proposing a parameter to linearly manipulate the stiffness of non-rigid objects. The stiffness parameter in most physical engines (e.g., Blender) is extremely non-linear. Thus, in the previous studies, the authors have to sample the parameters based on their experience to generate stimuli with perceptually plausible stiffness levels (Bi & Xiao, 2016; Bi, Jin, et al., 2018; Paulun et al., 2017; Schmid & Doerschner, 2018). In our method, the *α* value is linearly correlated with perceived stiffness (see Figure 6B), which allows us to easily create non-rigid objects that perceptually uniform in its stiffness.

## Conclusion

In conclusion, we discover that both intrinsic mechanical properties and optical properties affect stiffness perception of cloth when the stimuli are displayed as static images. In the video conditions, humans can partially discount the bias caused by optical appearances and exhibit higher sensitivity to stiffness. Analysis of optical flow fields shows that motion statistics associated with the pattern of dynamic deformation (i.e., movement uniformity) is diagnostic of stiffness of cloth. To further test how the patterns of dynamic deformation affect the inference of stiffness, we isolate the deformation information by removing the influence of optical properties and create dynamic dot stimuli from the 3D mesh of cloth animations. We propose a method to directly manipulate the pattern of dynamic deformation of the dot stimuli and demonstrate that changing movement uniformity of the dots can alter the perceived stiffness. We evaluate the robustness of this method by showing that it can be generalized to other scene set-ups, 3D model, and cloth physics. Finally, we confirm that the same six optical flow features (the mean and SD of divergence, gradient, and discrete Laplacian) are reliable image-based measurements to differentiate stiffness of non-rigid objects. Additionally, the linear combination of motion energy features is also diagnostic of the degree of stiffness of the dynamic dot stimuli. Together, our study demonstrates that manipulating patterns of dynamic deformation can elicit the impression of cloth with varying stiffness, suggesting the brain can infer mechanical properties from image cues related to dynamic image deformation.

## Supporting information

WindScene_HighSpeed_HighUniformity

WindScene_LowSpeed_LowUniformity

Exp1_RedGauze_WindScene

Exp1_Reference

Exp1_Felt_BallScene

Exp1_Cotton_BallScene

Exp1_Brocade_WindScene

Exp2_WindScene_alpha0.6

Exp2_WindScene_alpha1.2

Exp2_DrapeScene_alpha0.6

Exp2_DrapeScene_alpha1.4

Exp2_ConerScene_alpha0.4

Exp2_ConerScene_alpha1.4

Exp2_Cube_alpha0.4

Exp2_Cube_alpha1.2

